# A vagal sensory pathway for satiety suppression by intestinal fructose

**DOI:** 10.64898/2026.08.30.748157

**Authors:** Hikari Takeshima, Shigefumi Yokota, Mana Hatakeyama, Keisuke Ito, Kazunari Miyamichi, Hideki Enomoto, Takeshi Imai

## Abstract

Mammals primarily use glucose as an energy source, but they can also utilize other isocaloric monosaccharides, such as galactose and fructose. Since the gustatory system cannot distinguish between different sugars in ingested foods, it remains unclear whether and how mammals control the intake of different monosaccharides. Here we perform *in vivo* calcium imaging of the nodose ganglion in mice and find that intestinal glucose and fructose activate distinct populations of vagal sensory neurons. While glucose suppresses food intake in fed mice, fructose does not, resulting in increased calorie intake. However, after vagotomy, both glucose and fructose suppressed food intake, suggesting that vagal circuits activated by fructose facilitate feeding. This specialized vagal sensory pathway for fructose may partly explain the excess calorie intake induced by a fructose-rich diet.

## INTRODUCTION

Feeding is fundamental to acquiring the essential nutrients needed to maintain body homeostasis. Animals obtain carbohydrates, amino acids, and fats, as well as many other minerals and vitamins, from food. While glucose is a major source of energy for the cells in all animals, many animals can flexibly utilize other sugars, proteins, and fats as alternative energy sources. Animals can identify appropriate sets of nutrients in the foods using chemosensory systems. Additionally, they can detect the energy demands and nutrient availability in the body and control the feeding behavior using the hunger and satiety signals (Bruning and Fenselau, 2023). Although feeding is essential for survival, many people in modern society consume excessive amounts of food, which can lead to obesity and metabolic syndrome (Zuker, 2015; Blüher, 2019). With so many appetizing foods available nowadays, eating just the right amount of food is a challenge (Friedman, 2003).

Nutrients in the food are first detected by the gustatory system (Yarmolinsky et al., 2009; Liman et al., 2014). This system has five major modalities: sweet, umami, bitter, salty, and sour. Sweet taste, for example, represents various kinds of carbohydrates, such as glucose, fructose, and sucrose (Nelson et al., 2001). The five different taste modalities are detected by distinct types of taste receptor cells and trigger parallel neural circuits in the brain, known as labeled line circuits. After food intake, nutrients are also detected in the gut. Recent studies have demonstrated that enteroendocrine cells (EECs) in the intestine detect various nutrients in the gut and send the information not only to the humoral pathway but also to the neural pathway via vagal sensory neurons and the brainstem (Williams et al., 2016; Kaelberer et al., 2020; Liu and Bohórquez, 2022; Prescott and Liberles, 2022; Gruber et al., 2025). Some of these signals are also conveyed to the spinal cord (Goldstein et al., 2021; Kim et al., 2022). Vagal sensory neurons discriminate between some of the intestinal signals, such as glucose, fat, and water, in addition to stretch signals (Williams et al., 2016; Kaelberer et al., 2018; Bai et al., 2019; Tan et al., 2020; Ichiki et al., 2022; Li et al., 2022; Mcdougle et al., 2024). These signals control satiety and reward, contributing to food intake and preference (Han et al., 2018; Bai et al., 2019; Fernandes et al., 2020; Mcdougle et al., 2024). However, the diversity and the coding logic of the nutritional sensations by vagal sensory neurons are not fully understood.

In this study, we used *in vivo* calcium imaging to profile the responses of vagal sensory neurons in the nodose ganglion to various nutrients in the upper small intestine. Similar to gustatory systems, some of the vagal sensory neurons were responsive to particular nutrients. Furthermore, we found that glucose and fructose, which are perceived as sweet by the gustatory system, activate distinct sets of vagal sensory neurons that regulate distinct feeding behaviors in mice.

## RESULTS

### Comprehensive calcium imaging of vagal sensory neurons upon intestinal nutrient stimulation

The olfactory system has a divergent and combinatorial coding strategy, while the gustatory system has five distinct modalities and labeled line circuits (Firestein, 2001; Yarmolinsky et al., 2009; Liman et al., 2014). Vagal sensory neurons in the nodose ganglia consist of various transcriptional cell types, each of which receives distinct inputs from different visceral organs (Bai et al., 2019; Kupari et al., 2019; Zhao et al., 2022). Some of the vagal sensory neurons respond to chemicals in the intestinal lumen (Williams et al., 2016; Kaelberer et al., 2020). However, it remains to be determined how many types of nutrients are distinguished by these neurons.

We performed *in vivo* two-photon calcium imaging of the intact nodose ganglion using Avil-Cre; Ai162 and Phox2b-Cre; Ai38 mice. In these mice, GCaMP6s/GCaMP3 is expressed in all vagal sensory neurons (Zhou et al., 2010; Daigle et al., 2018; Putra et al., 2023). We exposed the nodose ganglion under the microscope, with their axons fully intact, because damage to the axons often impairs calcium responses. Since the imaging area was limited under these conditions, we used piezo z-drive to acquire time-lapse images at five different depths. We introduced nutrients from the pylorus of the stomach and perfused the upper small intestine (the proximal 12 cm of the small intestine corresponding to the duodenum and ileum) (Fig. 1A, Fig. S1A-C). When we slowly perfused the upper intestine (250 μL/min) with liquid food, we observed that calcium responses that were largely segregated from responses to gastric and intestinal distension, suggesting chemosensory responses (Fig. S1D, E). Using repeated intestinal glucose perfusion, we confirmed reproducible responses (Fig. S1F).

**Figure 1.**
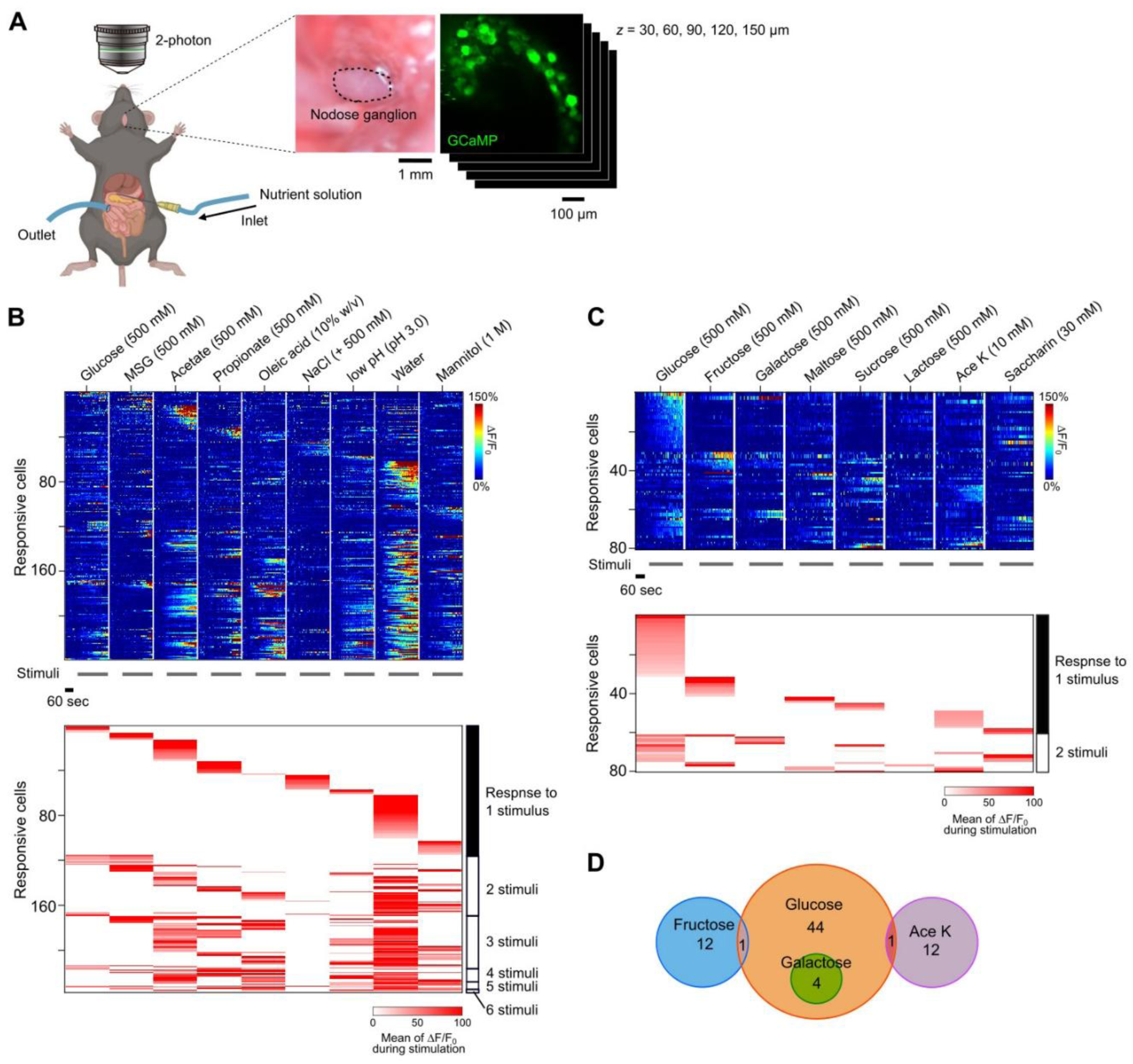
*In vivo* calcium imaging of the nodose ganglion reveals segregated circuits for intestinal nutrients. (A) Schematic diagram of *in vivo* two-photon calcium imaging of the nodose ganglion. We used Avil-Cre; Ai162 mice, in which GCaMP6s is expressed in vagal sensory neurons. Under anesthesia, the nodose ganglion was exposed and perfused with oxygenated Ringer’s solution. Using two-photon microscopy with a piezo z-drive, we imaged 5 different depths (30-150 μm, every 30 μm) at a rate 3.05 frame/sec for nutrient stimulation. The upper small intestine (∼12 cm) was perfused with PBS warmed to 37 °C at a rate of 250 μL/min. See also Fig. S1. (B) Calcium traces (top) and themean ΔF/F_0_ of responsive vagal sensory neurons to 9 different stimuli (239 responsive neurons out of 591 cells from 6 mice). Cells were sorted based on the number of stimuli to which they responded. The stimuli tested were glucose (500 mM), monosodium glutamate (MSG) (500 mM), acetate (500 mM), propionate (500 mM), oleic acid (500 mM), NaCl (PBS + 500 mM), low pH (pH 3.0), water, and 1 M mannitol (high osmolarity). The data show 239 responsive cells out of N = 591 cells identified (6 mice, 3 left nodose ganglia and 3 right nodose ganglia; note that non-responsive neurons with low basal GCaMP signals were not included). Comparison between left and right nodose ganglia are shown in Fig. S1H, I. (C) Calcium traces (top) and the mean ΔF/F_0_ of responsive vagal sensory neurons in response to 8 different sweeteners. The sweeteners tested were glucose (500 mM), fructose (500 mM), galactose (500 mM), maltose (500 mM), sucrose (500 mM), lactose (500 mM), acesulfame potassium (AceK) (10 mM), and saccharin (30 mM). The data show 87 responsive cells out of N = 399 cells (3 mice, left nodose ganglia). (D) Venn diagram of vagal sensory neurons that are responsive to glucose, fructose, galactose and AceK.

First, we performed calcium imaging of vagal sensory neurons while stimulating the upper intestine with different categories of nutrients and chemicals, including glucose (a sugar), glutamate (an amino acid), acetate and propionate (short-chain fatty acids), oleic acid (a long-chain fatty acids), NaCl (sodium), low pH, water, and mannitol (a high osmolality solution) (Fig. 1B). Of the 591 neurons recorded, 239 neurons responded to at least one of these stimuli. Of the responsive vagal sensory neurons, half were found to respond to just one type of stimulus. For example, responses to glucose, glutamate, fatty acids, sodium, and water were largely segregated. These results suggest segregated processing of multiple chemosensory signals, similarly to gustatory systems.

### Glucose and fructose activate distinct populations of vagal sensory neurons

Next, we examined whether vagal sensory neurons distinguish between different types of sugars. While animals consume various kinds of sugars, the major disaccharides utilized as energy sources are maltose, sucrose, and lactose. These disaccharides are digested into monosaccharides – namely, glucose, fructose, and galactose – at the intestinal brush border, and then absorbed by intestinal epithelial cells through various transporters. We examined responses to these monosaccharides, disaccharides, and artificial sweeteners (Fig. 1C). In the gustatory system, they are detected by the sweet receptor, Tas1r2/3 (Nelson et al., 2001). However, in nodose ganglion, very few neurons responded to all of them, suggesting that Tas1r2/3 are not the major receptor for the sugars in the intestinal lumen. Artificial sweeteners activated distinct populations of vagal sensory neurons, as previously reported (Buchanan et al., 2022). We also found that a substantial fraction of vagal neurons responds to fructose; these neurons almost did not overlap with the glucose-responsive population (Fig. 1D). This result suggests the existence of specialized sensory mechanisms for intestinal fructose.

### Transcriptional cell types of vagal sensory neurons activated by glucose and fructose

Next, we examined the transcriptional cell types of vagal sensory neurons that respond to glucose and fructose. Previous single-cell RNA sequencing studies have revealed 20-50 different transcriptional cell types among vagal sensory neurons (Bai et al., 2019; Kupari et al., 2019; Zhao et al., 2022). After intestinal perfusion with glucose and fructose, we identified activated neurons with antibodies against phosphorylated Erk (pErk), a neuronal activity marker (Fig. 2A, Fig. S2A-C) (Ji et al., 1999). Based on previous studies, we have chosen several marker genes that largely do not overlap and cover the major transcriptional cell types of vagal sensory neurons (Fig. 2C, Fig. S2D-F) (Zhao et al., 2022). We examined which marker genes are expressed in the pErk-positive neurons. We found that *Gpr65*^+^ and *Tmc3*^+^ neurons respond exclusively to fructose, whereas *Vip*^+^ neurons respond to either glucose or fructose (Fig. S3A-C). Further analyses revealed that *Ddc*/*Rbp4*, and *Glp1r*/*Uts2b* are more specific markers for *Tmc3*^+^ and *Vip*^+^ fructose-responsive neurons, respectively (Figs. 2C, D, Figs. S3D-I). *Uts2b*^+^ neurons responded to either glucose or fructose. However, *Gpr65*^+^ and *Rbp4*^+^ neurons only responded to fructose (Fig. 2D). Together, these results suggest the existence of at least three distinct transcriptional cell types responsive to intestinal fructose in the nodose ganglion.

**Figure 2.**
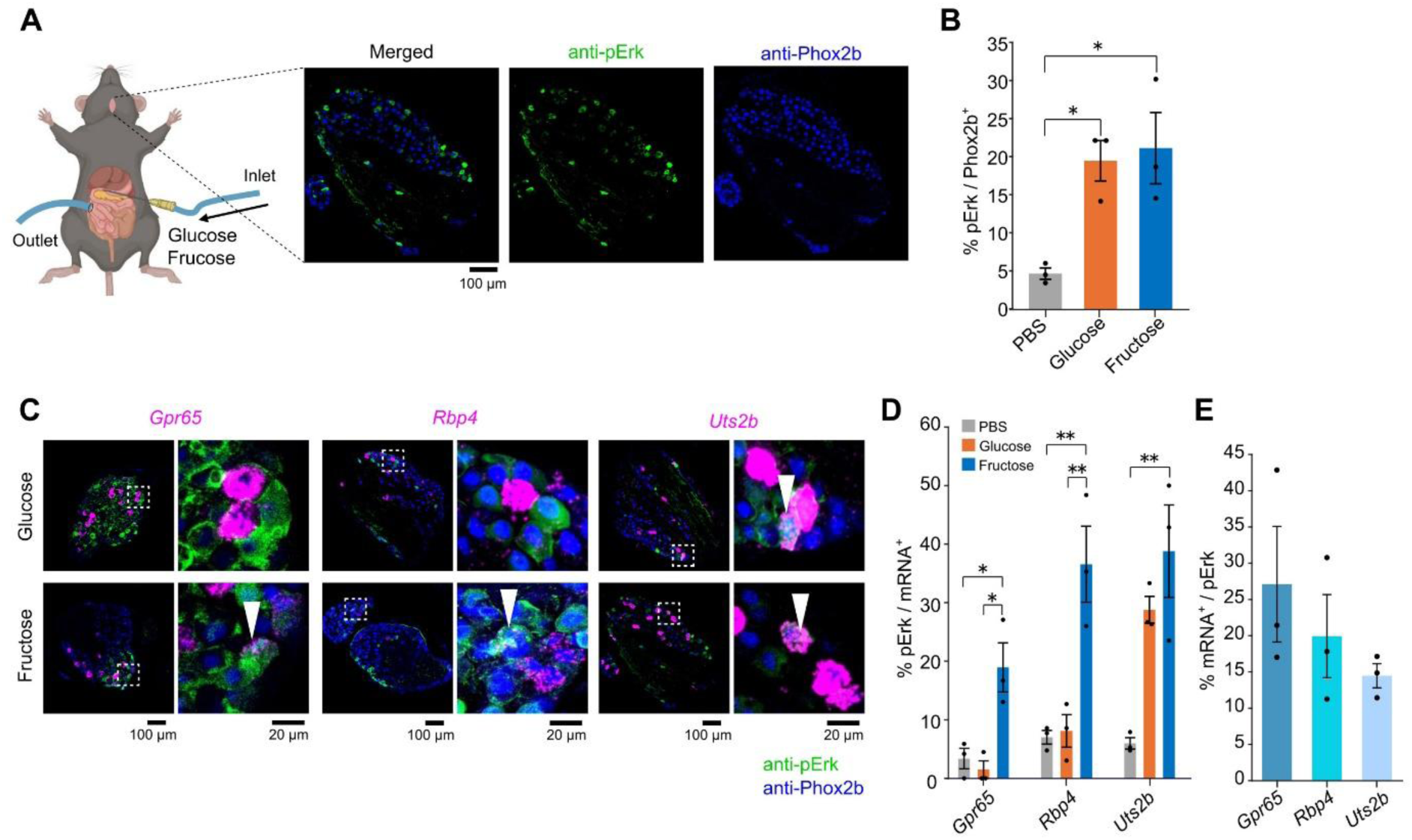
Transcriptional cell types of glucose-and fructose-responsive vagal sensory neurons. (A) Antibodies against phosphorylated Erk (pErk) label activated vagal sensory neurons in response to artificial perfusion of glucose/fructose into the upper intestine (∼12 cm). The transcriptional cell types of the activated neurons were identified using RNAscope. Anti-Phox2b labels all vagal sensory neurons. (B) Fraction of pErk-positive vagal sensory neurons after glucose/fructose perfusion of the upper intestine. N = 3 mice. *p*\* < 0.05 (one-way ANOVA followed by Tukey-Kramer multiple comparisons). (C) Specific gene markers for transcriptional cell types responsive to fructose. Previous studies have reported on the transcriptional cell types of the nodose ganglion using single-cell RNAseq. Starting from broadly expressed marker genes, we identified 3 specific clusters labeled with more specific marker genes (See also Fig. S3). The arrowheads indicate the double-positive neurons. (D) Quantification of pErk-positive neurons for specific cell types marked with RNAscope. *Uts2b*+ neurons responded to both glucose and fructose, while *Gpr65*+ and *Rbp4*+ neurons only responded to fructose. N = 3 mice. *p*\* < 0.05 (one-way ANOVA followed by Tukey-Kramer multiple comparisons). (E) *Gpr65*^+^, *Rbp4*^+^, and *Uts2b*^+^ neurons accounted for ∼60% of the fructose-responsive neurons. N = 3 mice.

A recent study reported that *Npy2r*^+^ vagal sensory neurons respond to glucose and fructose (McKnight et al., 2026). Indeed, *Glp1r*^+^/*Uts2b*^+^ neurons are known to express *Npy2r*. However, *Gpr65*^+^ and *Rbp4*^+^ neurons do not express *Npy2r* (Zhao et al., 2022). Thus, the *Gpr65*^+^ and *Rbp4*^+^ neurons likely comprise previously uncharacterized cell types that are specifically tuned to fructose but not to glucose in the upper intestine.

### Transcriptional cell types of cNTS neurons activated by glucose and fructose

Vagal sensory neurons project axons to the caudal part of the solitary tract (cNTS)(Cutsforth-Gregory and Benarroch, 2017). Neurons in the cNTS are classified into ∼20 transcriptional cell types, some of which have been demonstrated to control feeding behavior (Wang et al., 2025). We fed 10% glucose or fructose-containing water to starved mice and examined expression of c-Fos protein, a marker for neuronal activity (Fig. 3A-C). We then performed fluorescence *in situ* hybridization to identify which transcriptional cell types are activated by glucose and fructose. We found that *Gcg*^+^ cNTS neurons are activated by glucose but not fructose, consistent with a previous study using fiber photometry (Wang et al., 2025). Conversely, we found that *NPY*^+^ cNTS neurons are activated by fructose but not by glucose (Fig. 3D, E).

**Figure 3.**
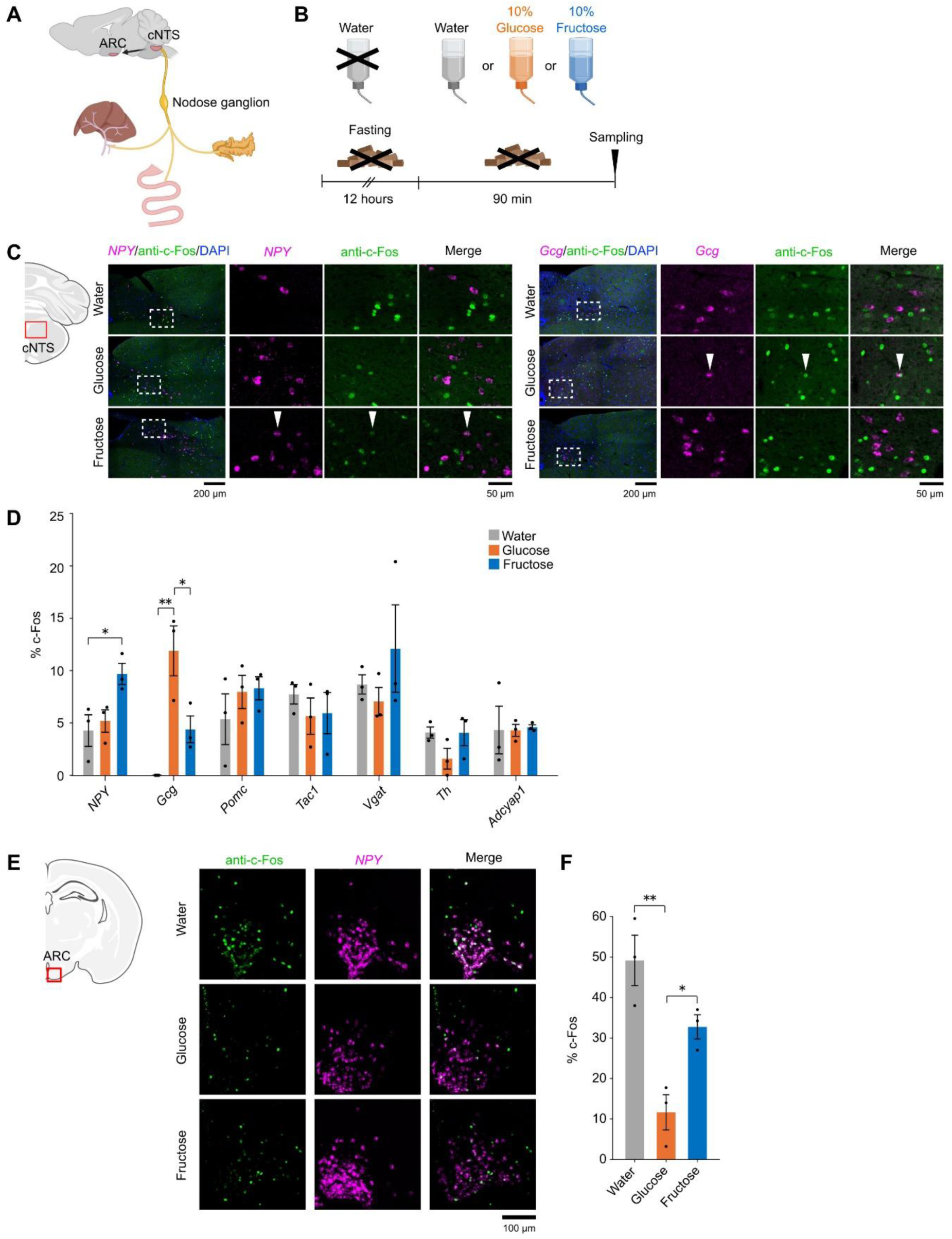
Glucose and fructose activate distinct feeding circuits in the brain. (A) Mapping glucose/fructose-activated neurons in the brain. We examined c-Fos expression in the caudal part of NTS (cNTS) and the arcuate nucleus (ARC). The cNTS receives inputs from vagus nerve, whereas the rostral part of NTS (rNTS) receives inputs from the gustatory system. Agrp neurons in the ARC play a key role in feeding regulation. Note that cNTS and ARC also receive non-vagal inputs. (B) Starved mice were allowed to drink water, 10% glucose water, or 10% fructose water for 90 min before c-Fos mapping. (C) c-Fos expression in the cNTS. Activated neurons were identified using specific marker genes identified in a previous study (Wang et al., 2025). *NPY* and *Gcg* genes are shown as examples. The arrowheads indicate the double-positive neurons. (D) Quantification of c-Fos expression in specific cNTS neurons. *Gcg*^+^ neurons were activated more strongly by glucose than by fructose. In contrast, *NPY*^+^ neurons were only activated by fructose intake. N = 3 mice. *p*\* < 0.05, *p*\* < 0.01 (One-way ANOVA followed by Tukey-Kramer multiple comparisons). (E) c-Fos expression in the ARC. While c-Fos was highly expressed in AgRP neurons (NPY^+^) in the water group, much lower expression was found in the glucose group, suggesting that glucose intake suppresses feeding. We observed a marginal reduction in c-Fos in the fructose group. (F) Quantification of c-Fos expression in AgRP neurons in the ARC. N = 3 mice. *p*\* < 0.05, *p*\* < 0.01 (One-way ANOVA followed by Tukey-Kramer multiple comparisons).

### Differential regulation of AgRP neurons

A previous study using Gq-DREADD experiments showed that *Gcg*^+^ cNTS neurons slightly reduce the food intake, while *NPY*^+^ cNTS neurons increase it (Chen et al., 2020; Wang et al., 2025). We, therefore, considered the possibility that glucose and fructose have distinct impacts on feeding regulation. It is known that a subset of cNTS neurons sends direct projections to the arcuate nucleus (ARC) of the hypothalamus, which plays a key role in regulating hunger and satiety (Aklan et al., 2020; de Morentin et al., 2024). We examined the regulation of agouti-related peptide (AgRP) neurons in the ARC, which positively control feeding (Deem et al., 2022; Bruning and Fenselau, 2023). Starved mice were fed 10% glucose or fructose for 90 min, after which c-Fos expression in AgRP neurons was examined. AgRP neurons were identified with anti-NPY antibodies. We found that c-Fos expression was reduced in glucose-fed animals. However, the fraction of c-Fos-positive neurons in the fructose-fed group was much higher than in the glucose group, and was comparable to that in the control (Fig. 3F). Our results with c-Fos staining are consistent with a recent study using fiber photometry (McKnight et al., 2026). These results suggest that glucose, but not fructose, suppresses feeding circuits in the brain.

### Regulation of feeding and satiety by glucose and fructose intake

Our results thus far have demonstrated that glucose and fructose activate distinct neuronal circuits in the brain. We then investigated how intake of these sugars modulate feeding behavior (Fig. 4A). Since mice typically consume more glucose water than fructose water (Sclafani et al., 1993), we presented the same amount (2 mL) of 10% glucose or fructose water, as well as adequate amount of water, during the 3-hour feeding assay. Under these conditions, the mice consumed all of the glucose or fructose solution. When we provided glucose and fructose water to starved mice, we did not observe any obvious differences in food intake (Fig. 4B). However, when we gave glucose or fructose water to fed mice, we observed differences between the glucose and fructose groups. Glucose-fed animals showed reduced food intake compared to the control group (water only), suggesting that glucose induces satiety signals. However, fructose feeding did not reduce the food intake.

**Figure 4.**
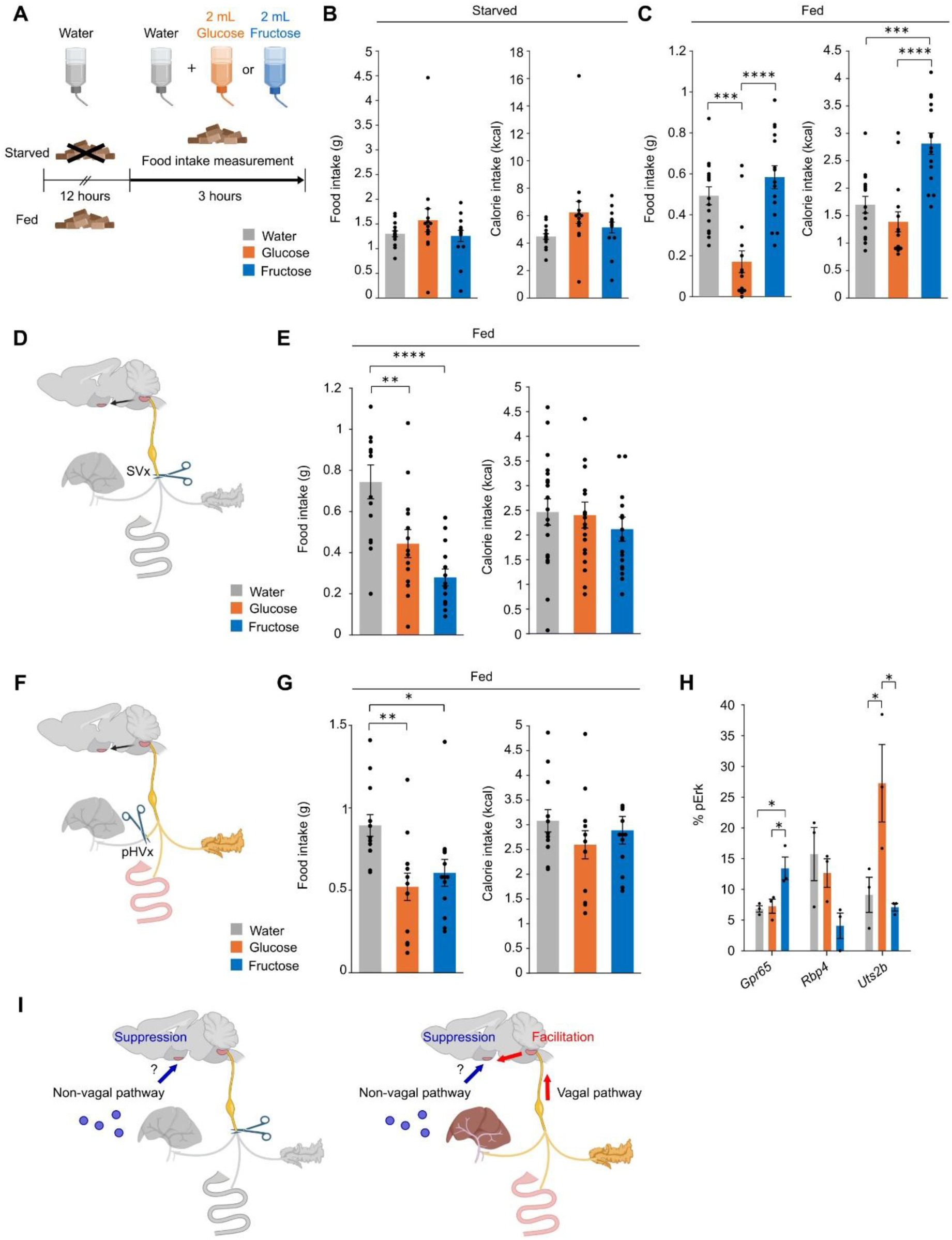
Distinct vagal modulation of feeding behavior by glucose and fructose. (A) Feeding assay. Starved (B) or fed (C) mice consumed 2 mL of either glucose (10%) or fructose (10%) water and had access to water during the feeding assay for 3 hours. The mice were habituated to the assay cage for 3 days beforehand. See also Fig. S4. (B) Food intake for fasted mice. Mice were fasted for 17 hours before the feeding assay. N = 15 mice per group. *p*\* < 0.01 (One-way ANOVA and Bonferroni multiple comparison test). (C) Food intake for fed mice. The mice had free access to normal foods at the test cage before the feeding assay. The fructose group showed higher calorie intake. N = 15 mice per group. *p*\*\*\*\* < 0.0001, *p*\*\*\* < 0.001 (One-way ANOVA and Bonferroni multiple comparison test). (D) Bilateral subdiaphragmatic vagotomy (SVx). The mice recovered for at least 5 days before the assay. Mice whose food intake did not return to normal after surgery were excluded from the feeding assay. (E) Food intake and calorie intake by SVx mice with glucose/fructose intake. Note that fructose suppresses food intake. N = 20 mice per group. *p*\*\*\*\* < 0.0001, *p*\*\* < 0.01 (One-way ANOVA and Bonferroni multiple comparison test). (F) Proper hepatic branch vagotomy (pHVx). A vagal nerve innervating the hepatic portal area was selectively transected. The mice were allowed to recover for at least 5 days before the feeding assay. Mice whose food intake did not return to normal after surgery were excluded from the feeding assay. (G) Food intake and calorie intake by pHVx mice with glucose/fructose intake. N = 13 mice per group. *p*\*\* < 0.01, *p*\* < 0.05 (One-way ANOVA and Bonferroni multiple comparison test). (H) pErk assay for upper intestinal glucose/fructose infusion in pHVx mice. Fructose-induced pErk signals in *Rbp4*-and *Uts2b*-positive nodose neurons were abolished by pHV. N = 3 mice. *p*\* < 0.05 (One-way ANOVA followed by Tukey-Kramer multiple comparisons). (I) A proposed model for fructose-induced feeding regulation. Fructose suppresses food intake via a non-vagal pathway but facilitates it via a vagal pathway.

Consequently, calorie intake in the fructose-fed group was higher than in the control and glucose-fed groups (Fig. 4C). We observed similar results for consecutive days (Fig. S4). Thus, glucose and fructose have distinct impacts on feeding and satiety regulation.

### Fructose facilitates feeding in fed mice via vagal sensory neurons

Next, we examined the role of the vagal sensory neurons in feeding regulation induced by glucose and fructose. We performed vagus nerve transection (vagotomy) and performed the feeding assay after recovery. Following subdiaphragmatic vagotomy (SVx), fructose water potently suppressed food intake in fed mice, similarly to glucose (Fig. 4D, E). These results suggest that fructose suppresses food intake via a non-vagal pathway, while facilitating it via vagal sensory neurons.

We further investigated which organ is responsible for the facilitation of food intake induced by fructose. When we transected the hepatic branch of the vagus nerve (HVx), feeding was suppressed by fructose intake (Fig. 4F, G). In the HVx mice, we observed reduction in pErk signals upon intestinal fructose stimulation in *Rbp4*^+^ vagal sensory neurons, but not in *Gpr65*^+^ neurons (Fig. 4H; see also Fig. 2D). Thus, the facilitation of feeding by fructose is at least in part mediated by *Rbp4*^+^ vagal sensory neurons that innervate the hepatic portal area (HPA).

## DISCUSSION

In this study, we found that intestinal glucose and fructose are detected by distinct populations of vagal sensory neurons. We also demonstrated that glucose and fructose differentially affect feeding and satiety regulation. A feeding assay is commonly performed on starved animals. In this study, we did not observe clear differences in food intake between animals fed with glucose and fructose in starved conditions (Fig. 4B), consistent with a previous study (McKnight et al., 2026). However, we found that glucose suppresses feeding while fructose does not in fed animals (Fig. 4C). In contrast, food intake was profoundly suppressed in vagotomized mice (Fig. 4E), suggesting that fructose suppresses food intake via non-vagal pathways while facilitating it via the vagal pathway. The non-vagal pathways may include humoral regulation and spinal cord pathways, as is known for glucose (Cummings and Overduin, 2007; Goldstein et al., 2021). Indeed, fructose is known to facilitate the secretion of insulin and some of gasterointestinal hormones (Kuhre et al., 2014). The state-dependent and transient nature of the action suggests that the fructose-dependent vagal pathway is not merely the induction of hunger. We assume that fructose suppresses satiety signals via vagal sensory neurons, thereby increasing total calorie intake (Fig. 4I).

We found that the hepatic branch of the vagus nerve is involved, at least in part, in the fructose-dependent facilitation of food intake. This pathway is most likely distinct from *Npy2r*^+^ vagal sensory neurons, which were recently reported to mediate fructose-induced suppression of AgRP neurons (i.e., feeding suppression) (McKnight et al., 2026). Currently, the fructose-specific sensing mechanism remains unknown. Luminal glucose is known to be transported to the intestinal epithelium by SGLT1 (Slc5a), while fructose is transported by GLUT5 (Slc2a5). However, we did not find any *Slc2a5*-positive and *Slc5a*-negative enteroendocrine cells in the duodenum (Fig. S5). One possible scenario is that fructose is detected by unknown cell surface receptors or intracellular machinery in EEC, which secrete a gastrointestinal hormone, and the vagus nerve in the hepatic portal area detects such hormones. It is also possible that the hepatic portal area directly detects fructose transported into the bloodstream.

Animals under energy needs become hungry and consume more food. They become satiated and stop eating based on the interoceptive feedback signals. Nutrients and stretch signals in the gastrointestinal tract induce the satiety and contribute to the tight feeding control (Cummings and Overduin, 2007; Bai et al., 2019; Goldstein et al., 2021). Ingested glucose is known to increase blood glucose levels and induce secretion of gasterointestinal hormones and insulin, which suppresses feeding (Coll et al., 2007; Bruning and Fenselau, 2023; Gruber et al., 2025). An excess amount of glucose is converted into glycogen in the liver and muscles. In contrast, fructose is converted to fat using ATP in the liver, helping animals to survive food shortages. Our results indicate that fructose transiently induce feeding in satiated mice via vagal sensory neurons, thereby increasing total calorie intake. Glucose is typically supplied from starch in the grains, which are relatively stable and available in the environment throughout the seasons. In contrast, fructose is most commonly found in fruits, which are only available during short periods of the year. It is therefore hypothesized that it is evolutionarily advantageous to consume fructose-rich foods more whenever possible and convert them to fat in preparation for food shortages (Johnson et al., 2023). The fructose-dependent vagal pathway identified in this study may play a key role in this process.

It remains unclear whether the same circuit mechanisms exist in humans. However, this vagal sensory circuit may explain why humans enjoy eating fructose-and sucrose-rich sweets even after they have felt full from the main dish. Notably, sucrose and fructose consumption has increased dramatically over the past 100 years. There is mounting epidemiological evidence suggesting an increase in fructose consumption leads to an increase in obesity and metabolic syndrome in modern society (Johnson et al., 2007; Tappy and Lê, 2010; Zuker, 2015; Blüher, 2019). It is therefore possible that the vagal pathway for intestinal fructose sensation will be an important target for controlling overeating and obesity.

## Acknowledgements

We thank Mariko Nishihara, Mako Shimizu, Jeon Dahyun, Emi Nozoe, and Satoko Hamatake for technical assistance; Stephen Liberles (Gpr65-Cre), Zachary Knight (Uts2b-Cre), Fan Wang (Advillin-Cre), and Hongkui Zeng (Ai9, Ai32, and Ai162) for mouse strains; Serina Yamada, Teppei Gogo, and Mitsue Hagihara for instruction of surgery and sharing reagents. We also appreciate technical support from The Research Support Center, Research Center for Human Disease Modeling, Kyushu University Graduate School of Medical Sciences, which is partially supported by the Mitsuaki Shiraishi Fund for BasicMedical Research. Some of the illustrations were created with BioRender.com.

## Funding

This work was supported by CREST program of Japan Science and Technology Agency (JST) (JPMJCR2021 to SY, KM, HE, and TI), Grants-in-Aids from the Japan Society for the Promotion of Science (JSPS), KAKENHI (JP21K19355, JP21H00205, JP21H05696, JP24H02308, and JP24H02312 to TI; JP22K11752, JP25K14891 to SY; JP25KJ1906 to HT), and the SPRING Grant of JST (JPMJSP2136 to HT). HT was a predoctoral research fellow (DC1) of JSPS.

## Author contributions

Conceptualization: TI; Methodology: HT, SY, KM; Validation: HT, TI; Formal analysis: HT; Investigation: HT (most of experiments), SY (*in situ* hybridization), MH; Resources: HT, SY, KI, HE, KM; Data curation: HT and TI; Writing – original draft: HT, TI; Writing – review & editing: all authors; Visualization: HT; Supervision and project management: SY, HE, KM, TI; Funding acquisition: SY, HE, KM, TI.

## Competing interests

The authors declare that they have no competing interests.

## Data and materials availability

All the image data will be deposited to SSBD:repository (https://ssbd.riken.jp/repository/). Source data for all the figures are in supplementary materials. No new program code was generated in this study.

## Methods

### Mice

All animal experiments were reviewed and approved by the Institutional Animal Care and Use Committee of Kyushu University (#A21-117, A23-073, and A25-185). The following mice were obtained from Jackson Laboratory and bred to C57BL/6J mice: Ai162D(TIT2L-GC6s-ICL-tTA2)-D (Ai162D) (JAX #031562) (Daigle et al., 2018), Ai38(RCL-GCaMP3) (Ai38) (JAX # 014538) (Zariwala et al., 2012). Advillin-Cre mice (Zhou et al., 2010) were obtained from RIKEN BRC (RRBRC10246) and were maintained on a C57BL/6N background. ePet-Cre was kindly provided by Dr. Deneris (Scott et al., 2005). Phox2b-Cre mice (Putra et al., 2023) were generated as described previously and maintained on a C57BL/6N background. C57BL/6N and C57BL/6J mice were purchased from Japan SLC. Mice were kept under a consistent 12-hour light/12-hour dark cycle (lights on at 8 a.m., and off at 8 p.m.), with an ambient temperature of 20-26°C and humidity of 40-70%. The mice always had access to water bottle and food, Rodent Diet CE-2 (CLEA Japan), unless otherwise noted. Both males and females were used for our imaging, histochemistry, and behavioral experiments, which produced consistent results. Ages are indicated in figure legends.

### *In vivo* calcium imaging of vagal sensory neurons in the nodose ganglion

We used GCaMP6s (Avil-Cre; Ai162) and GCaMP3 (Phox2b-Cre; Ai38) mice for calcium imaging. We could not obtain Phox2b-Cre; Ai162 mice, due to perinatal lethality. Data from GCaMP6s and GCaMP3 mice were analyzed separately. The mice were fasted for 12 hours and then anesthetized with ketamine (80 mg/kg, Daiichi-Sankyo) and xylazine (16 mg/kg, Bayer). During surgery, the depth of anesthesia was checked by pinching the hind paws with forceps. If the mouse showed a withdrawal reflex, an additional dose of anesthetics was administered. During imaging, an anesthesia unit for small animals (Univentor, 400 Anesthesia Unit) was used to administer isoflurane (FUJIFILM Wako, #099-06571) at a concentration of 0.5% at a rate of 700 mL/min to the mouse. Normal body temperature was maintained using a heat pad (NATSUME, #KN-475-3-35) during surgery, and a water-resistant silicone rubber sheet (Hakko Denki, #SBH2523) during imaging. The mouse was then fixed in the supine position with a surgical tape. Next, a 2 cm midline incision was made from the neck to the sternum. The submandibular and sublingual glands were detached to expose the trachea. The muscles on the surface of the trachea were split longitudinally with forceps to expose the tracheal cartilages. A small incision was made between the tracheal cartilages, and a 5 cm long silicone tube with an internal diameter of 0.5 mm (AS ONE, #1-596-01) was inserted into the lower part that connects to the lung. The tube insertion site was secured with medical-grade superglue (Aron Alpha A Sankyo, Daiichi Sankyo). Then, the trachea was displaced toward the 9 o’clock position using a surgical suture (NATSUME, F17-50B2). The digastric muscle was moved to the 1 o’clock position with a microhook (Muromachi Kikai, #10065-15) to expose the nodose ganglion. Additionally, the surrounding blood vessels were cauterized using a cautery kit (Bovie, #21601010). The nodose ganglion was perfused with oxygenated Ringer’s solution during imaging. Imaging of the nodose ganglion was performed with a two-photon microscopy (#MM201, Thorlabs), equipped with a 20x NA 0.5 water-immersion objective lens (Olympus/Evident, UMPLFLN20XW), and ThorImageLS software (Thorlabs). A 920-nm femtosecond laser (ALCOR 920-4 Xsight, SPARK LASERS) was used for excitation. During imaging, the objective lens was moved rapidly along the z-axis using a piezo actuator (PI, #P-725.2CD or nanoFAKTUR, #SFS-D20400) at 30-μm increments to a maximum depth of 150 μm. Images at 6 planes were acquired at 1.15 frame/sec for distension stimuli and at 3.05 frame/sec for nutrient stimuli. Both left and right nodose ganglia were imaged. In some experiments, we only present data for the left nodose ganglion.

Calcium imaging data were analyzed using ImageJ and MATLAB (Mathworks). Image Stabilizer plugin (https://imagej.net/plugins/image-stabilizer) was used for x-y motion correction. Region of interest (ROI) for vagal sensory neurons were manually determined. The ΔF signals were normalized to the mean intensity 90 sec prior to the stimulus onset (F_0_). The response amplitude was defined as the mean ΔF/F_0_ during the first 90 sec after the stimulus onset. Stimulus onset timing was estimated based on the pilot experiments with a dye solution (Fig. S1A-C). Neurons demonstrating response amplitudes above the mean + 3 SD of the baseline were defined as responsive neurons.

### Stimulation of the gastrointestinal tract for *in vivo* calcium imaging

Gastric distension was induced by inserting a 23G needle (Terumo, #NN-2332S) into the greater curvature of the stomach. Then, 1.5 mL of air was injected into the stomach using a 5 mL syringe (Terumo, #SS-05SZ). To perform intestinal distention, we ligated the pylorus of the stomach with surgical suture. Then, we inserted a 23G needle (Terumo, #NN-2332S) into the duodenum (∼1 cm away from the pylorus) and injected 2 mL of air using a 5 mL syringe (Terumo, #SS-05SZ).

To stimulate the upper small intestine (the duodenum and a part of the ileum), a 27G needle (Terumo, #NN-2719S) was inserted from the pylorus of the stomach into the duodenum to create an inlet for the solution. The intestine was cut ∼12 cm away from the pylorus and connected to a silicone tube with an internal diameter of 2 mm (AS ONE, #6-586-05) to create an outlet. The other end of the intestine was tied with suture to prevent the leakage of the intestinal contents. PBS was continuously perfused into the upper intestine at a rate of 250 μL/min using a peristaltic pump (M&S Instruments Inc., MINIPULS 3). During stimulation, PBS was replaced manually with a stimulus solution using a three-way stopcock (Nordson MEDICAL, #VXB1062). The stimulus valve was open for 90 sec, but the stimulus solution typically remained in the intestinal lumen for ∼3.5 min (Fig. S1A-C). After stimulation, the upper intestine was perfused with PBS for 15 min to flush out the stimuli. We used an in-line solution heater (Warnar Instruments, #SF-28) to heat the introduced solution to 37°C. Different stimuli were perfused in random sequences across trials.

The following stimuli were used: D-glucose (500 mM in PBS; Wako, #049-31165); monosodium glutamate (500 mM in PBS; adjusted to pH 7.3; Sigma, #G8415); acetate (500 mM in PBS; nacalai tesque, #00212-85); propionate (500 mM in PBS; TCI, #P0500); sodium chloride (500 mM in PBS; nacalai tesque, #31320-05); D-mannitol (1 M in PBS; Sigma, #M4125); D-fructose (500 mM in PBS; Wako, #127-02765); D-galactose (500 mM in PBS; Wako, #073-00031); maltose monohydrate (500 mM in PBS; Wako, #138-00611); sucrose (500 mM in PBS; Wako, #193-00025); lactose monohydrate (500 mM in PBS; Wako, #128-00095); acesulfame potassium (10 mM in PBS; Wako, #013-14102); saccharin (30 mM in PBS; Wako, #195-08662); and oleic acid (10% w/v in 0.1% xanthan gum and 0.05% Tween 80 in PBS, vortexed for 10 min prior to use; TCI, #O0011). For the oleic acid control, 0.1% xanthan gum (TCI, #X0048) and 0.05% Tween 80 (Sigma, #P1754) in PBS was used. To prepare a low pH (pH 3.0) solution, 36% HCl was added to PBS. For the liquid diet, we used Meibalance (Meiji, banana flavor, as an equivalent to Ensure Liquid, Abbott), diluted with water by half.

### Evaluating the timeline of intestinal perfusion

In our pilot experiments for calcium imaging, we perfused the upper intestine with 0.01% Fast Green (FUJIFILM Wako, #061-00031) in PBS. We collected the perfused solution in a 96-well plate (Thermo Scientific, #167008) every 30 sec at positions just before (inlet) and after (outlet) the upper intestine. The optical density (at 600 nm) of the solutions was recorded using a plate reader (Berthold, TriStar3). The optical density of PBS without Fast Green was then subtracted to determine the concentration of Fast Green. According to Figs. S1B, C, we estimate that the maximum concentration of the stimuli in the intestinal lumen is 60-100% of the perfused stimuli when the stimuli were perfused for 90 sec at a rate of 250 mL/min.

### Intestinal stimulation for pErk assay

Intestinal stimulation was performed as described for *in vivo* calcium imaging using C57BL/6N mice (2 to 4-months-old, males and females). The following stimuli were used: D-glucose (500 mM in PBS; Wako, 049-31165) and D-fructose (500 mM in PBS; Wako, #127-02765). Before stimulation, we exposed the nodose ganglion so that we could quickly dissect the nodose ganglia without unnecessary stimulation or damage. PBS was perfused for at least 30 min before stimulation with nutrients. Sampling was performed 20 min after the start of stimulation based on pilot experiments (Fig. S2E, F).

### Stimulation for c-Fos assay

Reliable c-Fos signals with low background were obtained through the oral administration of stimuli to awake animals. We used C57BL/6N mice (10 week-old, age-matched males). The mice were habituated to the experimental cage and bottles for 3 days prior to the experiments. According to a previous study (Tan et al., 2020), food and water were removed from the mouse cage for 12 hours prior to the experiments. During stimulation, the mice had access to a water bottle containing water, 10% glucose, or fructose (in MilliQ water) in the absence of food in the same cage. The animals were sacrificed 90 min later for c-Fos immunostaining.

We examined c-Fos expression in the rostral NTS (taste region), caudal NTS (visceral region), and ARC.

### *In situ* hybridization of vagal sensory neurons in the nodose ganglion

Mice were euthanized and perfused with PBS, followed by 4% PFA in PBS. The left nodose ganglion was dissected and post-fixed overnight at 4 °C in PFA in PBS. The samples were cryoprotected in 30% sucrose at 4°C for 1–2 days and embedded in OCT compound (Sakura). Frozen nodose ganglia were cut into 16-μm-thick coronal sections using a cryostat (Leica, #CM3050S). RNAscope (Advanced Cell Diagnostics) was used for *in situ* hybridization following the manufacturer’s instructions. Target probes were purchased from Advanced Cell Diagnostics. In double staining experiments, immunohistochemistry was performed after RNAscope. Briefly, after Amp6 of the RNAscope 2.5 HD Reagent Kit Brown (Advanced Cell Diagnostics, #322300), the sections were incubated with ClariTSA Fluorophore 520 (Advanced Cell Diagnostics, #323271). After washing in PBS, the sections were fixed with 4% PFA/PBS for 15 min. After washing in PBS, the sections were blocked (1% bovine serum albumin, 0.1% Triton X-100 in PBS) for 1h at room temperature. Next, the sections were incubated with primary antibodies overnight at 4 °C. The following primary antibodies were used: Human/Mouse Phox2B Antibody (1:500, R&D systems, AF4940), Phospho-p44/42 MAPK (Erk1/2) (Thr202/Tyr204) (pErk, 1:500, Cell Signaling, #9101), mCherry Goat Polyclonal Antibody (1:1000, ORIGENE, #AB0040-200). After washing 3 times for 15 min in PBS with 0.05% Tween20, the sections were stained with secondary antibodies for 30 min (pErk) or 2 h (others) at room temperature. Nuclei were stained with 0.1% DAPI. The following secondary antibodies were used: Alexa 488-conjugated donkey anti-goat IgG (1:500, ThermoFisher, #A11055), Alexa 647-conjugated donkey anti-rabbit IgG (1:500, ThermoFisher, #A31573). The samples were mounted on glass slides using ProLong Gold (Thermo Fisher, #P36930). Leica TCS SP8 (Leica) with a 20x objective (Leica HC PL APO 20×/0.75 IMH CORR C52, NA=0.75, WD=0.68mm) was used for confocal imaging.

We used the following RNAscope probes for the following mouse genes: *Calca* (#678771), *Ddc* (#318681), *Glp1r* (#418851), *Gpr65* (431431), *Tmc3* (#426301), *Trpa1* (#400211), *Trpv1* (#313331), *Uts2b* (#468331), *Vip* (#415961), *Rbp4* (#508501), *Slc2a5* (#508261-C2), and *Slc5a1* (#468881).

### *In situ* hybridization combined with immunohistochemistry of the brain (NTS and ARC)

After perfusion, the brains were immediately removed and post-fixed in the same fixative at 4°C for 24 h. The samples were then immersed in 25% sucrose in a 0.1 M phosphate buffer solution (pH 7.4). Once the brains had sunk, they were serially sectioned coronally at a thickness of 15 µm for the NTS and 30 µm for the ARC using a freezing microtome.

The sections were then washed in phosphate-buffered saline (PBS; pH 7.4) and pre-hybridized for 1 h at 60°C in a hybridization buffer containing Denhardt’s solution (0.02% bovine serum albumin, 0.02% Ficoll, and 0.02% polyvinylpyrrolidone; Eppendorf), 0.1% Tween 20 (ICN Biomedicals Inc.), 0.5 mg/ml yeast tRNA (EMD Millipore Corp.), 5× saline-sodium citrate (SSC; 150 mM NaCl, 15 mM sodium citrate, pH 7.0), 5% dextran sulfate (EMD Millipore Corp.), and 50% formamide (Fujifilm). The sections were then hybridized for 16 h at 60°C in the same buffer containing digoxigenin (DIG)-and fluorescein isothiocyanate (FITC)-labeled riboprobes. The sections were washed twice in 2× SSC containing 50% formamide. Then, they were washed in 1× SSC containing 50% formamide and 0.2× SSC containing 50% formamide, both at 60°C for 1 h. Finally, the sections were rinsed in Tris-buffered saline (TBS; pH 7.3).

Antisense probes were designed for the coding sequences of the following genes (along with their NCBI accession#): *Adcyap1* (NM_009625.2, nt. 989-1850, DIG), *Cck* (NM_031161.1, nt. 2-608, DIG), *Gcg* (NM_008100.2, nt. 135-947, FITC), *Npy* (NM_023456.2, nt. 15-489, FITC), *Pomc* (NM_008895.2, nt. 31-945, FITC), *Tac1* (NM_009311.1, nt. 95-994, DIG), *Vgat* (NM_009508.3, nt. 887-1805, DIG).

Riboprobe detection was performed using the fluorochromized tyramide-glucose oxidase method described previously (Yamauchi et al., 2022). Since both DIG-and FITC-labeled riboprobes were used, the first riboprobe was detected, followed by the second. Signals were visualized using Cy3-tyramide, Cy5-tyramide, or biotin-tyramide followed by fluorophore-conjugated streptavidin.

For the first detection, the sections were incubated overnight in TBS containing 0.3% Triton X-100, 1% blocking reagent (Roche), and either a peroxidase-conjugated sheep anti-DIG antibody (1:500; Roche) or a peroxidase-conjugated sheep anti-FITC antibody (1:500; Roche). The sections were then incubated for 30 min in TBS containing 20 µM Cy3-or Cy5-conjugated tyramide (AAT Bioquest), or 10 µM biotin-tyramide (Tocris), along with 3 µg/mL glucose oxidase, 2 mg/mL D-glucose, and 1% bovine serum albumin. After the first riboprobe was detected, the sections were incubated in TBS containing 2% sodium azide for 4 h to inactivate the peroxidase conjugated to the antibody. Following washes in TBS, the sections were incubated overnight in TBS with the peroxidase-conjugated antibody for the remaining riboprobe. The sections were then reacted with a different tyramide substrate than that used in the first round of detection.

After *in situ* hybridization, the sections were incubated overnight in PBS containing 0.3% Triton X-100 and rabbit anti-c-Fos antibody (1:1000; Abcam). Then, the sections were incubated with Alexa Fluor 488 Plus-conjugated donkey anti-rabbit IgG (1:1000; Thermo Fisher Scientific Inc) for 3 h. In some experiments, the sections were incubated overnight with a mouse anti-tyrosine hydroxylase antibody (1:2500; ImmunoStar), followed by incubation with Alexa Fluor 405 Plus-or Alexa Fluor 647 Plus-conjugated donkey anti-mouse IgG (1:1000; Thermo Fisher Scientific Inc) for 3 h.

The sections were mounted onto gelatin-coated glass slides and coverslipped with VECTASHIELD Plus Antifade Mounting Medium (Vector Laboratories). Images were acquired using a spinning disk confocal microscope (Andor, Dragonfly 200) mounted on an inverted microscope (Nikon, Eclipse Ti2) and controlled by Fusion software (Andor).

### Feeding assay combined with sugar water intake

The feeding assay was performed for C57BL/6J mice (2 month-old males) using a Multi-feeder (SHINFACTORY, #MS-3F), which was located on the home cage. The mice were habituated to the feeder for 3 consecutive days prior to the feeding assay. During this time, the mice were fed with the feeder 3 hours a day. The weight of the Multi-feeder containing food pellets (CE2, CLEA Japan) was measured before and after the experiment to determine the food intake. During the feeding assay, the mice were provided with an adequate amount of water in a bottle and 2 mL of 10% glucose/fructose water in another. The starved groups were deprived of foods for 17 hours prior to the feeding experiments while having access to an adequate amount of water. All the feeding experiments were conducted during the dark phase. Data were excluded from the analysis if mice failed to feed from the feeder during the last day of habituation (food intake < 0.1 g in 3 hours). Food contained 3.45 kcal/g and 10% glucose/fructose water contained 0.4 kcal/mL.

### Vagal nerve transection

The mice were anesthetized with anesthetized with ketamine (80 mg/kg, Daiichi-Sankyo) and xylazine (16 mg/kg, Bayer). An incision was made along the midline of the abdomen. For subdiaphragmatic vagotomy (SVx), the stomach was exposed and the anterior vagal trunk, which runs along the abdominal esophagus, was severed with fine forceps. The stomach was then turned around, and the posterior vagal trunk was cut in the same way. For the proper hepatic branch vagotomy (pHVx), the left lobe of the liver was first displaced to expose the portal vein. Then, the vagal nerve running alongside the portal vein was cut with fine forceps. After the denervation, the animals were allowed to recover for at least 5 days, until their food intake returned to normal. Mice whose food intake did not return to normal after surgery were excluded from the feeding assay.

## QUANTIFICATION AND STATISTICAL ANALYSIS

### Image data processing and quantification

RNAscope images of the nodose ganglion were analyzed using ImageJ and MATLAB. Phox2b immunostaining was used to identify vagal sensory neurons. Cells containing four or more RNA puncta within the soma were defined as mRNA-positive according to previous literatures. Over 500 Phox2b neurons in four sections were analyzed per animal per condition. The proportion of pErk-positive cells among mRNA-positive neurons was quantified manually with ImageJ. *In situ* hybridization images of the NTS were analyzed similarly using ImageJ and MATLAB. The percentage of mRNA-positive neurons among c-Fos-positive neurons was quantified manually using ImageJ.

## Statistical analysis

GraphPad Prism7 and MATLAB (MathWorks) were used for the statistical analyses. Statistical tests used in the analyses are described in figure legends.

## Supplementary Information

**Figure S1.**
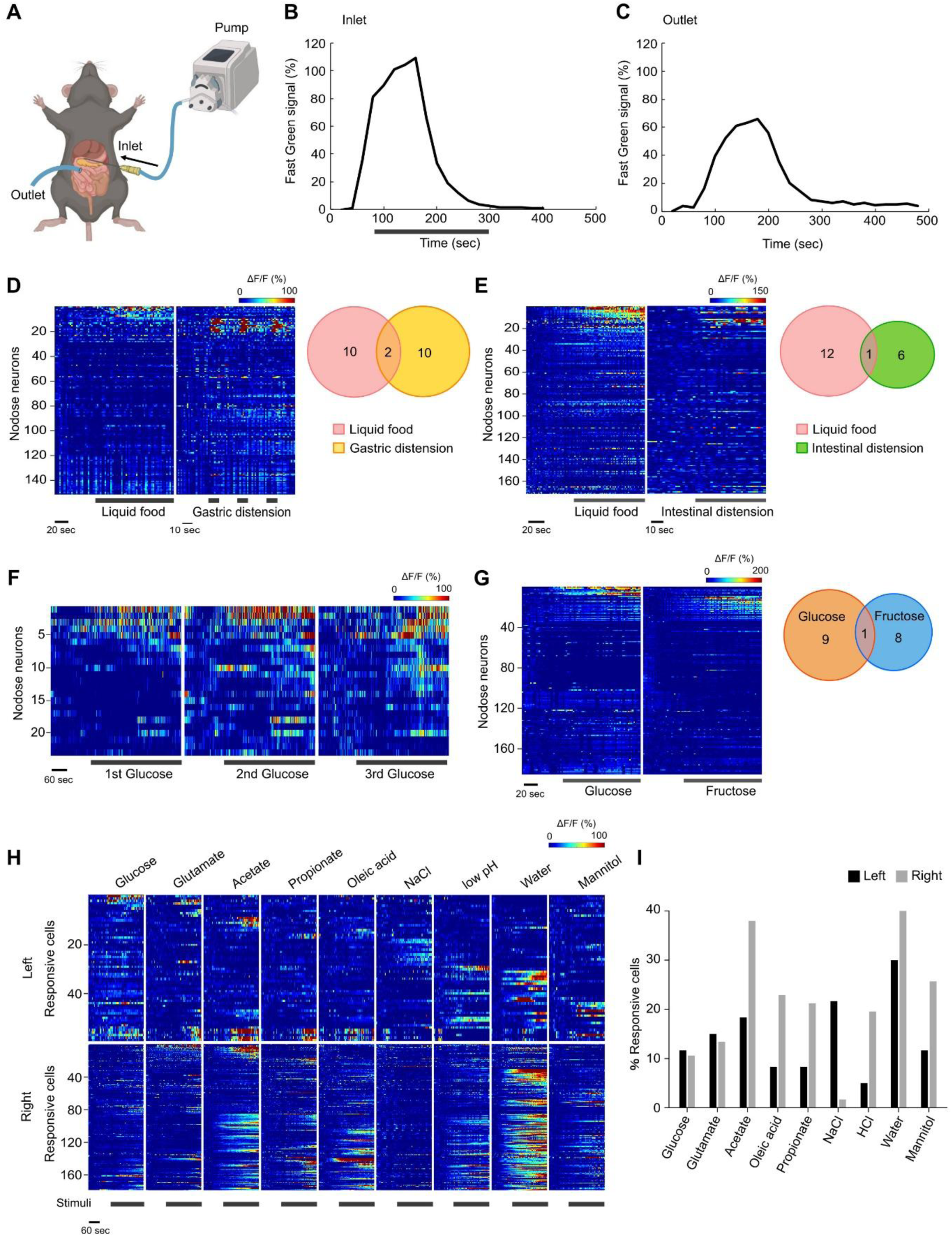
*In vivo* calcium imaging of vagal sensory neurons. (A) Estimation of stimulation timing using Fast Green dye. Using the *in vivo* imaging setup, we determined the dye concentrations at the inlet and outlet positions. (B) Temporal changes in Fast Green concentration at the inlet position. The concentration of the original Fast Green solution was defined as 100%. Throughout this study, the stimulation timing was defined as the 90 sec period during which the dye concentration at the inlet position was greater than 80% (shown by the gray horizontal bar). The end of stimulation was defined as 300 sec, when the dye concentration returned to baseline (shown by the gray horizontal bar). (C) Temporal changes in Fast Green concentration at the outlet position. (D) A comparison of liquid food (50% Meibalance) versus gastric distension stimuli (1.5 mL of air). The left nodose ganglion was imaged (data from 2 mice). The Venn diagram shows little overlap in responses between these two stimuli. GCaMP3 mice (Phox2b-Cre; Ai38) were used. (E) A comparison of liquid food (50% Meibalance) versus intestinal distension stimuli (2 mL of air). The left nodose ganglion was imaged (data from 2 mice). The Venn diagram shows little overlap in the responses to these two stimuli. GCaMP3 mice (Phox2b-Cre; Ai38) were used. (F) Reproducibility of nutrient stimulation. Glucose stimulation (500 mM in PBS) was performed three times on the same animals, demonstrating reproducible responses. The left nodose ganglion was imaged (data from 2 mice). GCaMP3 mice (Phox2b-Cre; Ai38) were used. (G) Responses to glucose and fructose in GCaMP3 mice (Phox2b-Cre; Ai38) (data from 2 mice). (H) Calcium responses of the left versus right nodose ganglion. The original data are the same as in Fig. 1B. The data are from 3 left nodose ganglia and 3 right nodose ganglia. GCaMP6s mice (Avil-Cre; Ai162) were used. The lateralization found for some nutrients may reflect differences in the innervation patterns of the left and right nodose ganglia, as well as the differential distribution of nutrient sensors (Bai et al., 2022; de Araujo et al., 2023). (I) Quantification of responsive cells in the left and right nodose ganglia. The fraction of responsive cells differed for some stimuli between the left and right ganglia.

**Figure S2.**
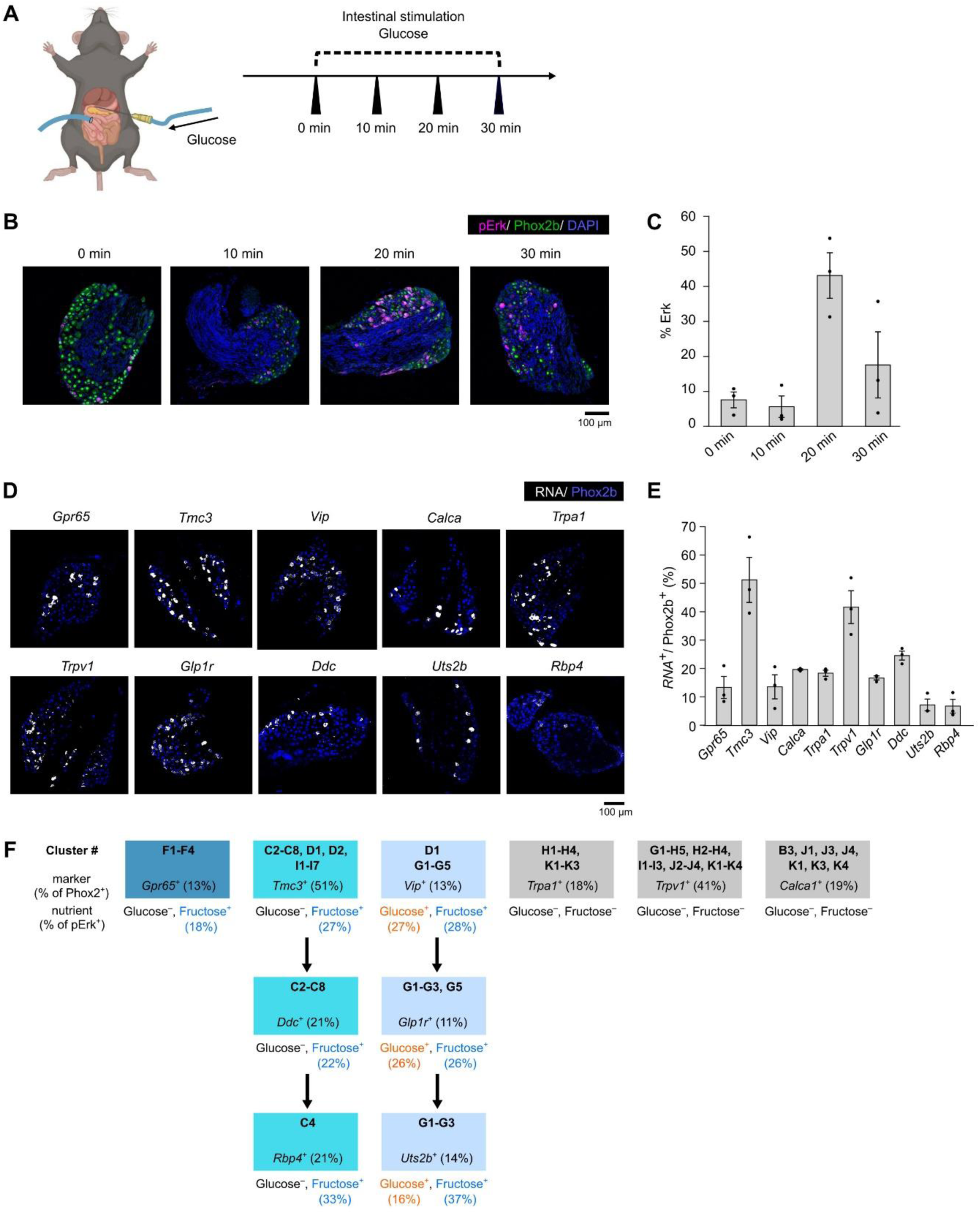
Activity mapping in the nodose ganglion. (A) Optimization of activity mapping with anti-pErk immunostaining. Anesthemized mice were perfused with glucose (500 mM in PBS) for up to 30 min. The nodose ganglion was then isolated and immunostained with anti-pErk antibodies. (B) Immunostaining of the left nodose ganglion after glucose perfusion to the upper intestine. (C) Quantification of pErk-positive vagal sensory neurons in the left nodose ganglia. N = 3 mice. Based on this pilot experiment, we sacrificed the mice 20 min after the starting nutrient perfusion in this study. (D) *In situ* hybridization (RNAscope) of the nodose ganglion with each of the marker genes. (E) Fraction of vagal sensory neurons positive for each or the marker genes. N = 3 mice. (F) Identification of specific marker genes for fructose-responsive vagal sensory neurons. The cluster numbers of transcriptional cell types are from Zhao et al. (Zhao et al., 2022). See Fig. S3 for the raw data in each step.

**Figure S3.**
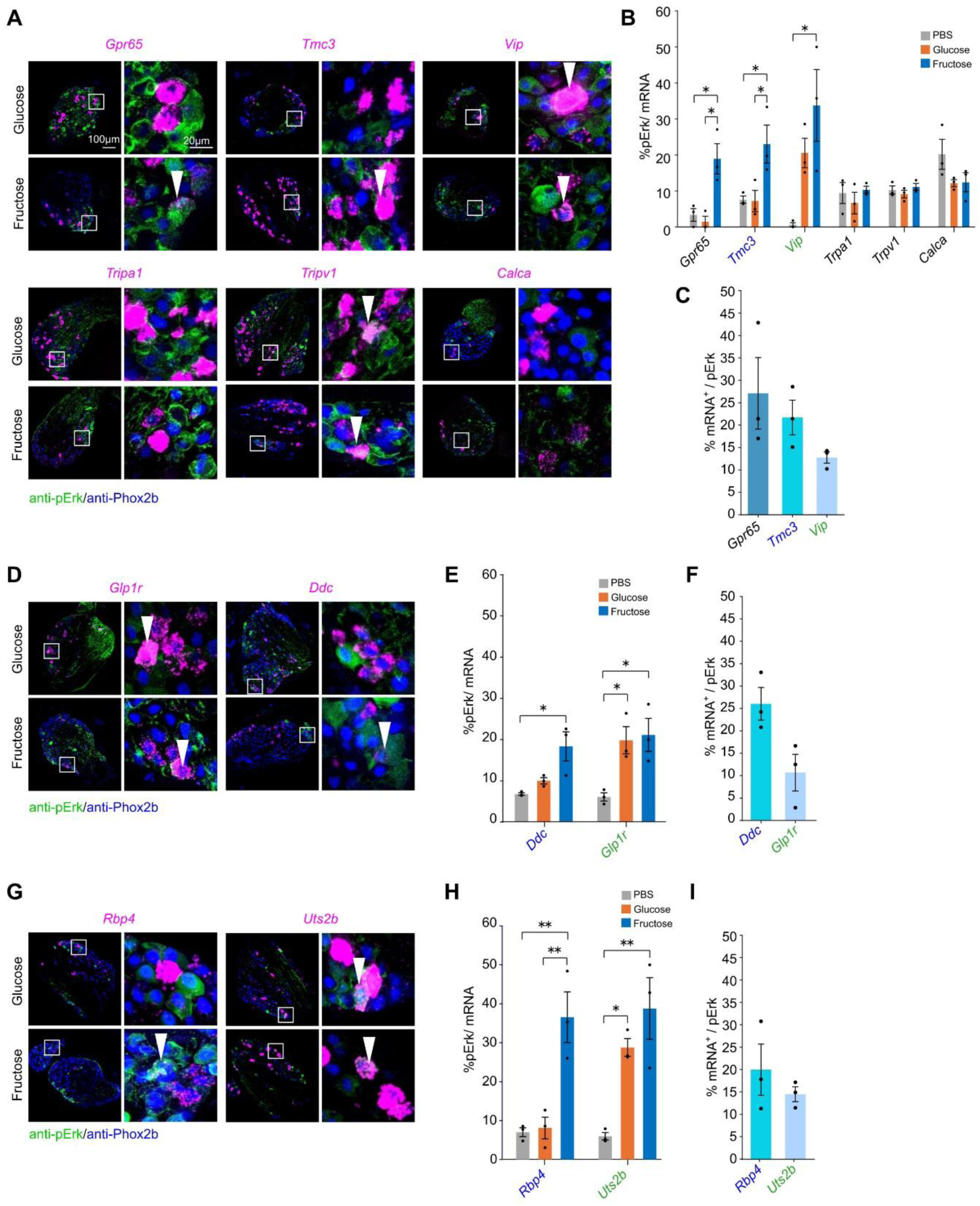
Transcriptional cell types for glucose-and fructose-responsive neurons. (A) In the first round, we have selected *Gpr65*, *Tmc3*, *Vip*, *Trpa1*, *Trpv1*, and *Calca* as markers for gut-innervating vagal sensory neurons, based on a previous study (Zhao et al., 2022). Responses to glucose or fructose were found in *Gpr65*^+^, *Tmc3*^+^, and *Vip*^+^ neurons. The arrowheads indicate the double-positive neurons. (B) Fraction of pErk^+^ neurons within each marker-positive neuron population. N = 3 mice. *p*\* < 0.05 (One-way ANOVA followed by Tukey-Kramer multiple comparisons). (C) Quantification of marker-positive neurons among pErk^+^ neurons. N = 3 mice. (D-F) In the second round, we identified *Gpr1r* and *Ddc* as more specific markers. N = 3 mice. *p*\* < 0.05 (One-way ANOVA followed by Tukey-Kramer multiple comparisons). (G-I) In the third round, we identified *Rbp4* and *Uts2b* as more specific markers. N = 3 mice. *p*\*\* < 0.01, *p*\* < 0.05 (One-way ANOVA followed by Tukey-Kramer multiple comparisons).

**Figure S4.**
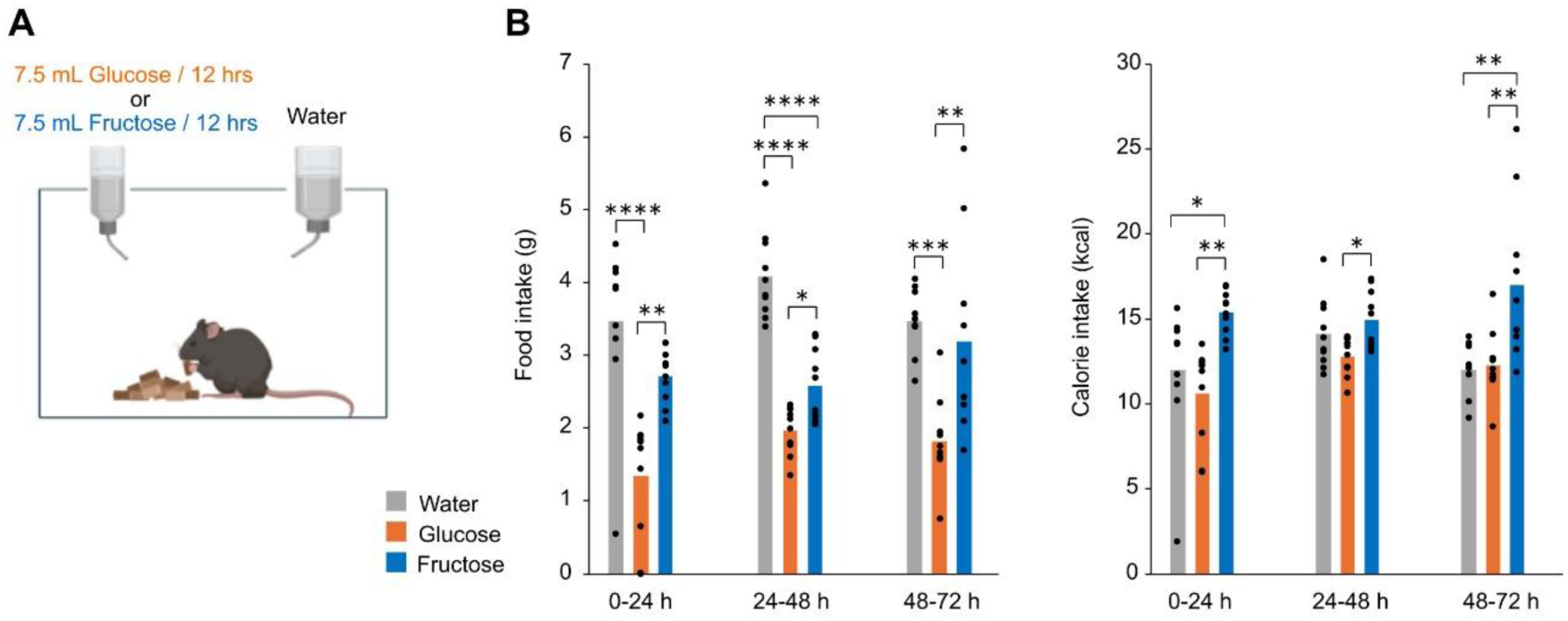
Feeding measurements for 3 days. (A) Daily feeding assay. 7.5 mL of Glucose or Fructose water (10%) was provided every 12 hours. Water was freely accessible. (B) Food intake and calory intake every 24 hours. N = 10 mice. *p*\*\*\*\* < 0.0001, *p*\*\* < 0.01, *p*\* < 0.05 (One-way ANOVA and Bonferroni multiple comparison test).

**Figure S5.**
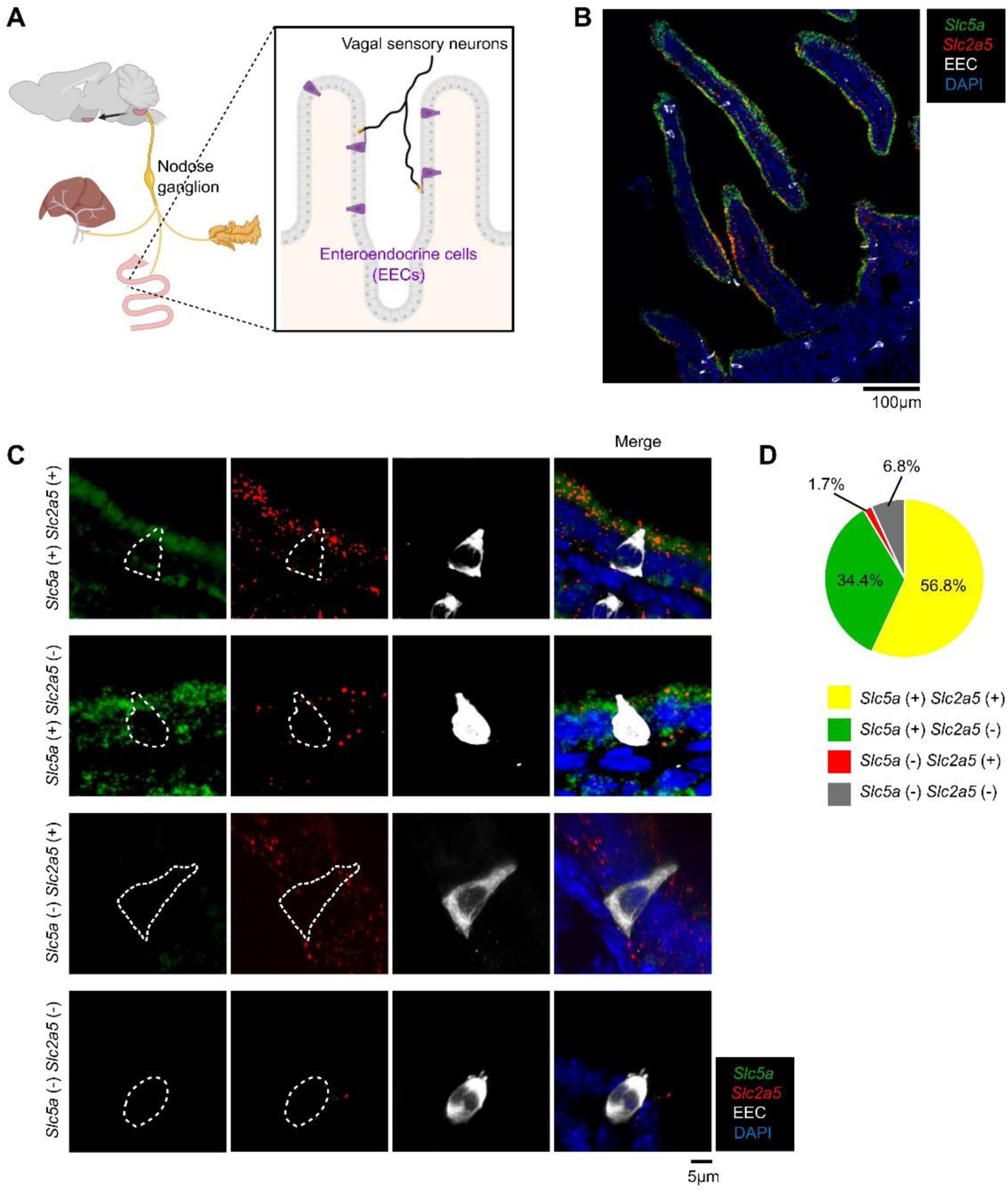
Expression of glucose and fructose transporters in the gut. (A) Schematic diagram of the small intestine. Enteroendocrine cells are sparsely scattered in the intestinal epithelium. (B, C) RNAscope double staining for *Slc5a* (coding for SGLT1) and *Slc2a5* (coding for GLUT5) was performed in the duodenum. The EEC was labeled with ePet-Cre (ePet-Cre; Ai9), which is a pan-EEC-specific Cre line (Bai et al., 2022). (D) Quantification of RNAscope. N = 48 cells from 1 mouse.

## Notes

### Competing Interest Statement

The authors have declared no competing interest.

